# Mycorrhizal status impacts the genetic architecture of mineral accumulation in field grown maize (*Zea mays* ssp. *mays* L.)

**DOI:** 10.1101/2022.12.12.520122

**Authors:** Meng Li, Sergio Perez-Limón, M. Rosario Ramírez-Flores, Benjamín Barrales-Gamez, Marco Antonio Meraz-Mercado, Gregory Ziegler, Ivan Baxter, Víctor Olalde-Portugal, Ruairidh J. H. Sawers

## Abstract

Arbuscular mycorrhizal fungi (AMF) establish symbioses with major crop species, providing their hosts with greater access to mineral nutrients and promoting tolerance to heavy metal toxicity. There is considerable interest in AMF as biofertilizers and for their potential in breeding for greater nutrient efficiency and stress tolerance. However, it remains a challenge to estimate the nutritional benefits of AMF in the field, in part due to a lack of suitable AMF-free controls. Here we evaluated the impact of AMF on the concentration of 20 elements in the leaves and grain of field grown maize using a custom genetic mapping population in which half of the families carry the AMF-incompatibility mutation *castor*. By comparing AMF-compatible and AMF-incompatible families, we confirmed the benefits of AMF in increasing the concentration of essential mineral nutrients (*e*.*g*., P, Zn, and Cu) and reducing the concentration of toxic elements (*e*.*g*., Cd and As) in a medium-input subtropical field. We characterised the genetic architecture of element concentration using quantitative trait mapping and identified loci that were specific to AMF-compatible or AMF-incompatible families, consistent with their respective involvement in mycorrhizal or direct nutrient uptake. Patterns of element covariance changed depending on AMF status and could be used to predict variation in mycorrhizal colonisation. We comment on the potential of AMF to drive genotype-specific differences in the host ionome across fields and to impact the alignment of biofortification breeding targets. Our results highlight the benefits of AMF in improving plant access to micronutrients while protecting from heavy metals, and indicate the potential benefits of considering AMF in biofortification programs.

## INTRODUCTION

Arbuscular mycorrhizal fungi (AMF) are a widely distributed group of soil fungi that form mutualistic associations with more than 80% of terrestrial plant species including the major staple crops (Wang & Qiu, 2006). One of the major benefits of AMF to hosts is enhanced phosphorus (P) acquisition through an extensive root-external hyphal network that accounts for up to 90% of the total P uptake (van der Heijden *et al*., 2015). Arbuscular mycorrhizal (AM) symbioses also play important roles in promoting the uptake of other macro- and micronutrients such as N, S, Zn, and Cu, and mitigating the toxicity of heavy metals, such as Cd, As, and Pb (Lehmann & Rillig, 2015; Ruytinx *et al*., 2020). Previous studies have reported that the nutritional benefits of AMF extend to the edible tissues of various crops (Pellegrino & Bedini, 2014; Pepe *et al*., 2022; Gupta *et al*., 2022), suggesting the potential for enhanced food quality and biofortification. Given the nutritional benefits provided by AMF, there is a growing interest in breeding for more AMF-effective crops as a contribution to enhanced plant nutrient use efficiency (Cobb *et al*., 2021; Thirkell *et al*., 2022). However, although AM symbioses have been shown to be beneficial in many greenhouse studies, there is a lack of support at the field scale due to environmental heterogeneity and the lack of non-mycorrhizal controls (Ryan & Graham, 2018). In addition, the low heritability of AMF colonisation and its inconsistent relationship with plant growth and nutrient content suggest the need to search for more reliable breeding targets for AM fungal benefits (Leiser *et al*., 2016; Feldmann, 2009).

Field evaluation of the nutritional benefits of AM symbioses is challenging, as it is difficult to manipulate AM communities without changing other variables. Field inoculation can be complicated by the presence of native fungal communities and is not cost-effective at a large scale in many cropping systems (Ortas, 2012; Salomon *et al*., 2022). Applying fungicides to experimentally reduce the abundance of native AMF has limited effect and may impact non-target fungal species, disrupting other biotic interactions and complicating the interpretation of the results (Buysens *et al*., 2015). Other strategies using agricultural management practices to manipulate AM fungal communities, such as plant rotation with non-host species or tillage, are confounded by other factors, such as soil water and nutrient availability (Ryan & Graham, 2018). As a result, reported estimates of host nutritional responses to AMF at the field scale vary widely (Lehmann *et al*., 2014; Lehmann & Rillig, 2015). Comparing AMF-incompatible mutant plants to their AMF-compatible wild-type siblings is an attractive complementary strategy to evaluate the impact of the AM symbiosis under field conditions (Watts-Williams & Cavagnaro, 2015; Bowles *et al*., 2016; Ryan & Graham, 2018; Ramírez-Flores *et al*., 2020).

Traditionally, studies of plant nutrition focused on different elements in isolation. However, the accumulation of nutrients in plants is strongly influenced by the interactions among elements, through soil chemistry, physiological disruption caused deficiency or toxicity, and crosstalk among plant nutrient signalling pathways (Baxter, 2015). Profiling the ionome, the complete set of mineral nutrients and trace elements in an organism (Lahner *et* al., 2003), provides an opportunity to comprehensively understand element accumulation and patterns of interaction. AMF can improve the content of some nutrient elements, such as P, by direct transportation through fungal networks (Chiu & Paszkowski, 2019). AMF can also impact host nutrient status indirectly by altering root architecture or as a consequence of alleviating primary deficiencies (Ramírez-Flores *et al*., 2019). Indeed, the concentration of a number of elements can be significantly changed in response to AMF inoculation (Hart & Forsythe, 2012; Ramírez-Flores *et al*., 2017). It is possible that the acquisition, translocation, and homeostasis of one element may be affected by the capacity of plants to acquire other nutrients directly or indirectly through AMF. The interplay between elements and AMF can be further complicated by soil nutrient availability. As reviewed in a meta-analysis, AMF increased tissue Cu, Mn, Fe and S levels under P deficiency, but not when P was sufficient (Watts-Williams & Cavagnaro, 2014). Given the impact of complex interactions among elements and AM symbiosis, it is informative to consider the ionome as a whole to fully understand the nutritional benefits of AM symbiosis under field conditions.

Plant roots can take up nutrients directly from the soil (the *direct* pathway) or acquire them via symbiosis with AMF (the *AMF* pathway; (Smith *et al*., 2003)). In the case of P, specific members of the PHT1 phosphate transporter family have been identified to be involved in the direct and AMF pathways, their expression and accumulation patterns responding to colonisation by AMF (Bucher, 2007; Chiu & Paszkowski, 2019; Salvioli di Fossalunga & Novero, 2019). For example, MtPT4, OsPT11 and ZmPT6 accumulate predominantly in the periarbuscular membrane surrounding arbuscules in *Medicago truncatula*, rice and maize, respectively (Harrison *et al*., 2002; Sawers *et al*., 2017; Paszkowski *et al*., 2002; Willmann *et al*., 2013; Fabiańska *et al*., 2020). Conversely, transporters implicated in direct P uptake are repressed in response to AMF (Bucher, 2007; De Vita *et al*., 2018). Although currently less well defined, analogous direct and AMF pathways appear to also function in N uptake, driven by specific accumulation of distinct ammonium transporters (Hui *et al*., 2022; Koegel *et al*., 2013). Further transporter families (for example, the ZIP Zn transporters (Nguyen *et al*., 2019); or SULTR sulphate transporters (Casieri *et al*., 2012)) have been reported to show gene-specific patterns of induction or repression during AM symbiosis, suggesting the concept of direct and AMF uptake to extend across the ionome, with the potential for AMF to influence crosstalk during nutrient homeostasis (Xie *et al*., 2019). The accumulation of nutrient transporter genes involved in direct and AMF pathway can vary among plant genotypes, *e*.*g*., phosphate transporter genes in maize (Sawers *et al*., 2017), indicating that host genetic factors may have a large impact on mycorrhizal-mediated nutrient uptake of plants.

Here we evaluated the ionome of a bi-parental maize genetic mapping population in which half the families carry the mutation *castor* and are incompatible with AMF. By comparing AMF-compatible and AMF-incompatible families, we estimated the importance of AMF on the accumulation of 20 elements in the field. We characterised the genetic architecture of host element accumulation in response to AMF and identified genetic regions associated with element concentration variation in AMF-compatible and incompatible families. Furthermore, we tested the capacity of ionome profiles to predict mycorrhizal status and assessed the potential of breeding AMF-effective hosts to achieve biofortification targets.

## RESULTS

### The *castor* mutation impacted the leaf and grain ionome of field grown plants

To assess the impact of AM symbioses on host mineral element accumulation in the field, we used inductively coupled plasma mass spectrometry (ICP-MS) to evaluate the concentrations of 20 elements in leaf and grain samples from a previously described custom mapping population (see Materials and Methods and (Ramírez-Flores *et al*., 2020)) grown in a subtropical, medium input, field in central-western Mexico. The population was generated from the cross between W22 (temperate) and CML312 (subtropical) maize inbred lines and contained both AMF-compatible (AMF-C; 75 families) and AMF-incompatible (AMF-I; 66 families) families to allow estimation of the overall impact of AMF and the characterization of genetic architecture underlying ionome differences among the families. Among the 20 tested elements, 17 in leaves and 9 in grain differed significantly in concentration between AMF-C and AMF-I families, including both essential mineral nutrients and toxic heavy metals (Fig. 1; Table 1). Element response to AMF (*Mycorrhizal Response* - MR; defined here as the percentage increase in AMF-C compared with AMF-I) ranged from −36% to 116% in leaves and from −25% to 65% in grain (Fig. 1A). Zn and Cu concentrations showed the highest positive response in both leaves (116% and 71%, respectively) and grain (50% and 65%, respectively). The concentrations of potential toxic elements, such as As (−36%) and Mn (−24%) in leaves, and As (−20%) and Cd (−23%) in the grain, were significantly reduced in AMF-C plants, consistent with previous reports (Lehmann & Rillig, 2015; Ruytinx *et al*., 2020). We conducted Principal Component (PC) analyses, and AMF-C and AMF-I ionomes were separated by the first PCs in both leaf and grain, which captures AM effect driven by Zn, Cu, and Fe (Supplementary Fig. S1). Although not the most responsive element, leaf P concentration was significantly increased in AMF-C compared to AMF-I families (8.5%). Grain P concentration did not change significantly with AMF status. In our study field, both AMF-C and AMF-I families maintained an average P concentration within a typical critical range (Gagnon *et al*., 2020), indicating that P limitation was not a major driver of previously reported differences in plant performance (Ramírez-Flores *et al*., 2020).

**Table 1.** Element concentrations (p.p.m) in leaves and grain of mycorrhizal compatible (AMF-C) and incompatible (AMF-I) families. Means and standard deviation (S.D.) were calculated from 75 and 66 genotypes for AMF-C and AMF-I, respectively. The significance of mean differences between two families were tested using the Wilcoxon test with p-values adjusted (FDR adjustment) based on the number of elements. Note: *: p < 0.05; **: p < 0.01; ***: p < 0.001; NS: not significant. Broad sense heritability (H^2^) of the whole mapping population was estimated using the *bwardr::Cullis_H2* function for R.

|  | Leaf |  |  |  |  |  | Grain |  |  |  |  |  |
| --- | --- | --- | --- | --- | --- | --- | --- | --- | --- | --- | --- | --- |
| | AMF-C | | AMF-I | | p | $H^2$ | AMF-C | | AMF-I | | p | $H^2$ |
| Element | Mean | S.D. | Mean | S.D. |  |  | Mean | S.D. | Mean | S.D. |  |  |
| Al | 11.17 | 3.29 | 11.61 | 3.33 | NS | 0.51 | 0.57 | 0.16 | 0.55 | 0.16 | NS | 0 |
| As | 0.04 | 0.02 | 0.07 | 0.02 | *** | 0.65 | 0.01 | 0.002 | 0.01 | 0.002 | *** | 0.26 |
| B | 18.61 | 3.66 | 16.36 | 4.53 | ** | 0.21 | 0.54 | 0.11 | 0.56 | 0.12 | NS | 0 |
| Ca | 6279.35 | 1682.39 | 7040.19 | 1653.11 | * | 0.64 | 83.57 | 23.97 | 90.79 | 29.70 | NS | 0.28 |
| Cd | 0.03 | 0.01 | 0.03 | 0.01 | NS | 0.73 | 0.004 | 0.002 | 0.005 | 0.002 | * | 0.58 |
| Co | 0.06 | 0.02 | 0.06 | 0.02 | NS | 0.53 | 0.004 | 0.001 | 0.004 | 0.001 | NS | 0.13 |
| Cu | 5.57 | 0.94 | 3.25 | 0.90 | *** | 0.69 | 1.71 | 0.30 | 1.03 | 0.37 | *** | 0.4 |
| Fe | 75.70 | 14.32 | 117.97 | 31.65 | *** | 0.62 | 20.52 | 2.81 | 27.50 | 6.25 | *** | 0.2 |
| K | 30672.02 | 3391.24 | 33142.20 | 4283.85 | ** | 0.61 | 5921.39 | 659.15 | 6016.12 | 635.67 | NS | 0.14 |
| Mg | 2826.21 | 488.60 | 3237.95 | 615.78 | *** | 0.71 | 1813.79 | 168.99 | 1849.13 | 185.64 | NS | 0.21 |
| Mn | 148.62 | 37.70 | 196.77 | 46.80 | *** | 0.55 | 5.47 | 1.02 | 6.15 | 1.12 | *** | 0.39 |
| Mo | 0.40 | 0.07 | 0.36 | 0.06 | ** | 0.31 | 0.07 | 0.07 | 0.05 | 0.07 | NS | 0.03 |
| Na | 10.26 | 4.16 | 8.24 | 3.15 | ** | 0.1 | 0.45 | 0.09 | 0.49 | 0.10 | NS | 0 |
| Ni | 0.04 | 0.01 | 0.03 | 0.01 | ** | 0.17 | 0.03 | 0.01 | 0.03 | 0.01 | * | 0.46 |
| P | 2510.42 | 242.64 | 2313.33 | 322.66 | *** | 0.63 | 2662.30 | 281.83 | 2770.97 | 348.63 | NS | 0.14 |
| Rb | 7.15 | 1.28 | 6.68 | 1.23 | * | 0.4 | 1.74 | 0.43 | 1.57 | 0.44 | NS | 0 |
| S | 3790.03 | 275.70 | 3683.31 | 318.64 | * | 0.51 | 1970.28 | 244.41 | 2019.22 | 198.12 | NS | 0.23 |
| Se | 0.02 | 0.01 | 0.02 | 0.00 | ** | 0 | 0.01 | 0.003 | 0.01 | 0.003 | ** | 0 |
| Sr | 27.42 | 9.06 | 30.15 | 7.95 | * | 0.66 | 0.42 | 0.16 | 0.52 | 0.18 | * | 0.32 |
| Zn | 40.89 | 8.67 | 18.88 | 8.61 | *** | 0.44 | 19.58 | 3.16 | 13.08 | 3.46 | *** | 0.25 |

**Figure 1.**
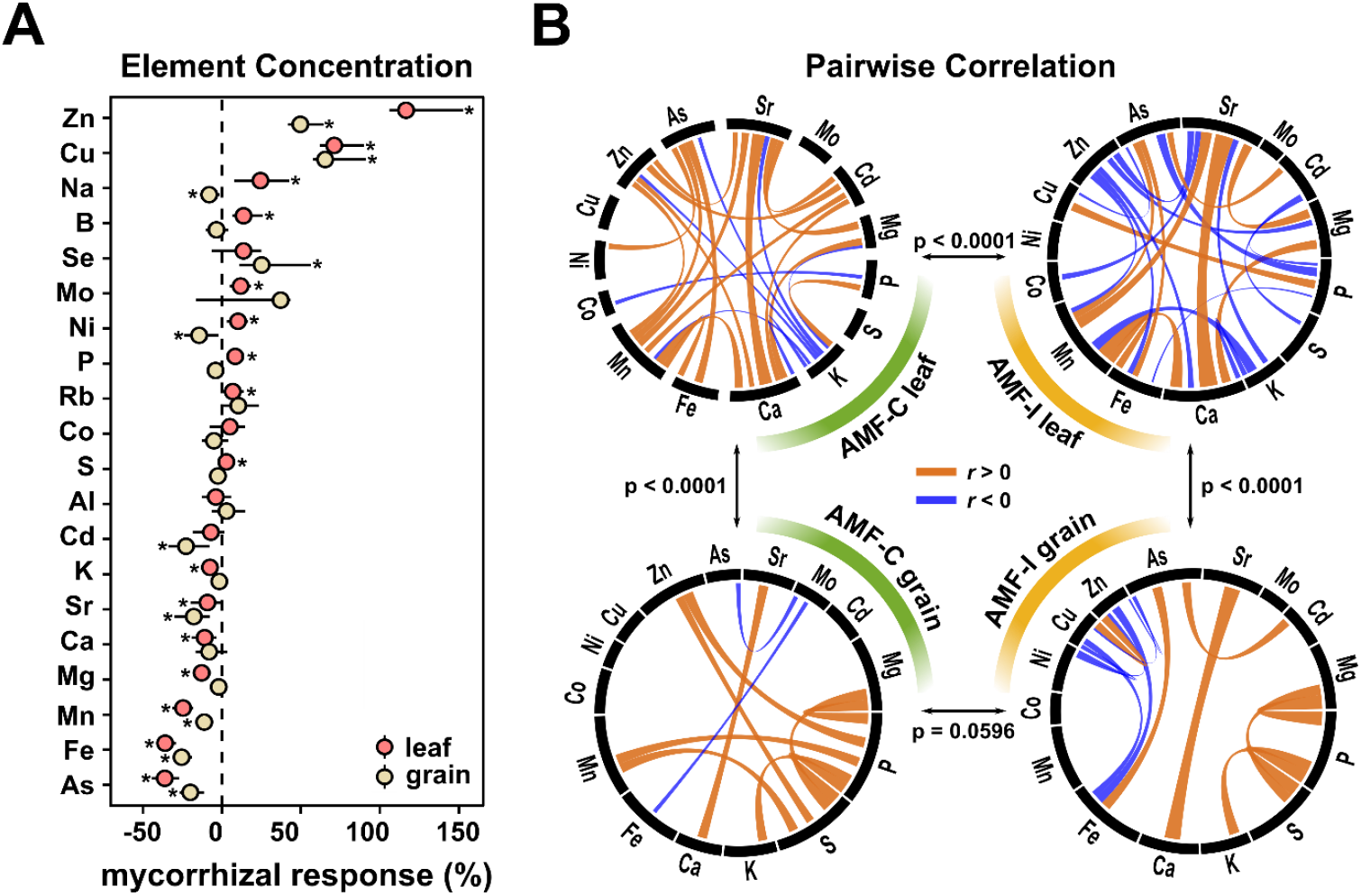
Mycorrhizal colonisation affects element concentrations and correlations in field grown maize. A) Mycorrhizal responses of element concentration in leaf and grain. Mycorrhizal response of each element was calculated as the mean difference between element concentration in compatible (AMF-C) and incompatible families (AMF-I) divided by AMF-I. Error bars represent 95% confidence intervals. B) Pairwise correlations of element concentration in leaf (upper) and grain (bottom) between AMF-C (left) and AMF-I (right) families. Significant Spearman correlations (p < 0.05) shown with positive correlations in orange and negative correlations in blue. The width of links represents the absolute value of correlation coefficients. Differences in correlation matrices were tested using the Chi-square test.

In addition to impacting individual elements, AM status affected patterns of covariance among elements. AMF-C and AMF-I families were more clearly distinguished by leaves than grain as shown by the magnitude of MR associated with individual elements (Fig. 1A). This pattern was supported by significant differences between pairwise correlation matrices for element concentrations in the leaves of AMF-C and AMF-I families (p < 0.001; Fig. 1B). For AMF-C and AMF-I leaves (Fig. 1B), 11 of the significant correlations were independent of AM status (*e*.*g*., the positive correlations between Sr and Ca, or As and Fe). In contrast, the sign of three correlations changed with AM status, indicating a strong effect of AMF on covariance (*e*.*g*., the positive correlation between Zn and the three elements Sr, Ca and Mn in AMF-C became negative in AMF-I). Ten significant correlations were specific to AMF-C (*e*.*g*., the positive correlations between K and P) and 10 to AMF-I (*e*.*g*., the positive correlation of P and Cu) families. The grain element correlation matrix, unlike that of leaves, was not significantly different between AMF-C and AMF-I families (p = 0.0596; Fig. 1B), although only five pairwise correlations were shared across AM status of a total of 11 and 14 significant correlations in AMF-C and AMF-I families, respectively (Fig. 1B). We also conducted Factor Analysis to reduce the dimension of element accumulation patterns and extracted the latent variables that share common trends in differentiating AMF-C and AMF-I for downstream analyses (Supplementary Fig. S2). In summary, AM status substantially affected patterns of covariance among elements, with more than half of the significant correlations changing depending on AMF status in leaf and grain.

### The genetic architecture of element accumulation was impacted by AMF status

Having observed a significant general effect of AMF status on the plant ionome, we proceeded to use Quantitative Trait Loci (QTL) linkage mapping to evaluate genetic differences between W22 and CML312 hosts and AMF × QTL effects. As discussed previously, we equated AMF-C and AMF-I specific QTL with mycorrhizal *benefit* and *dependence*, respectively (Ramírez-Flores *et al*., 2020; Sawers *et al*., 2010). In the context of the ionome, AMF × QTL effects may also reflect a distinction between AMF and direct plant uptake pathways (Smith *et al*., 2003). To identify AMF × QTL effects, we ran a series of QTL models: 1) all families without taking AMF status into account; 2) all families with AMF status as an additive covariate; 3) all families with AMF status as an interactive covariate; 4) AMF-I families alone; 5) AMF-C families alone. Model 2 eliminates the main effect of AMF status, but does not allow for the effect of W22 and CML312 alleles to differ with AMF status (*i*.*e*., AMF × QTL effects are not modelled). AMF × QTL effects are captured explicitly by model 3, and by comparing the results of models 4 and 5. We considered leaf and grain element concentrations as separate traits in our analysis.

Across all models, we detected a total of 33 element QTL, considering QTL for the same element detected in the same genetic bin in different models as the same. When QTL for different elements colocalized to the same bin, we report them as distinct loci, although it is likely they represent a single pleiotropic genetic variant. Similarly, we report QTL detected for leaf and grain data as distinct. QTL were annotated by element, chromosome, genetic bin, and tissue (Fig. 2). Thirteen QTL were detected in the grain and 20 in the leaves. In a single instance (qCd 2.05), a QTL for the same element is co-localised in leaf and grain, presumably representing a single causal locus. Fifteen QTL were detected specifically in AMF-C families but not AMF-I families, and, conserversely, 5 QTL were detected in AMF-I but not AMF-C families. Six of these AMF status specific QTL were further supported as examples of AMF × QTL interaction by Model 3. A further QTL (qAs_gr_8.09) was identified uniquely by Model 3. The AMF × QTL effects we detected were all *conditional* in nature (*i*.*e*., specific to AMF-C or AMF-I families). We did not find evidence of any QTL showing strong, but opposing, allelic effects between AMF-C and AMF-I families (*antagonistic pleiotropy*. See *e*.*g*., (Ramírez-Flores *et al*., 2020)). For the AMF-C specific QTL qCd_lf/gr_2.05 and qNi_gr_9.01, the effect was sufficiently strong that they were supported by Models 1 and 2, but not Model 3 (*i*.*e*., the interactive model did not present marked improvement over the additive model). Globally, the parental allelic effect of detected QTLs varies depending on the element and tissue type (Fig. 2). For example, for QTLs detected in bin 4.08 for leaf in AMF-C families, the genotypes that carry the CML312 allele tend to accumulate less Ca, Fe and As, but more Rb. For qCd_lf/gr_2.05, the allelic effect is conserved across grain and leaf in the AMF-C families, where the genotypes with W22 allele tend to accumulate less Cd than genotypes with the CML312 allele. In contrast, for qNi_gr_9.01, the AMF-C with the W22 allele at this QTL tends to accumulate Ni in the grain.

**Figure 2.**
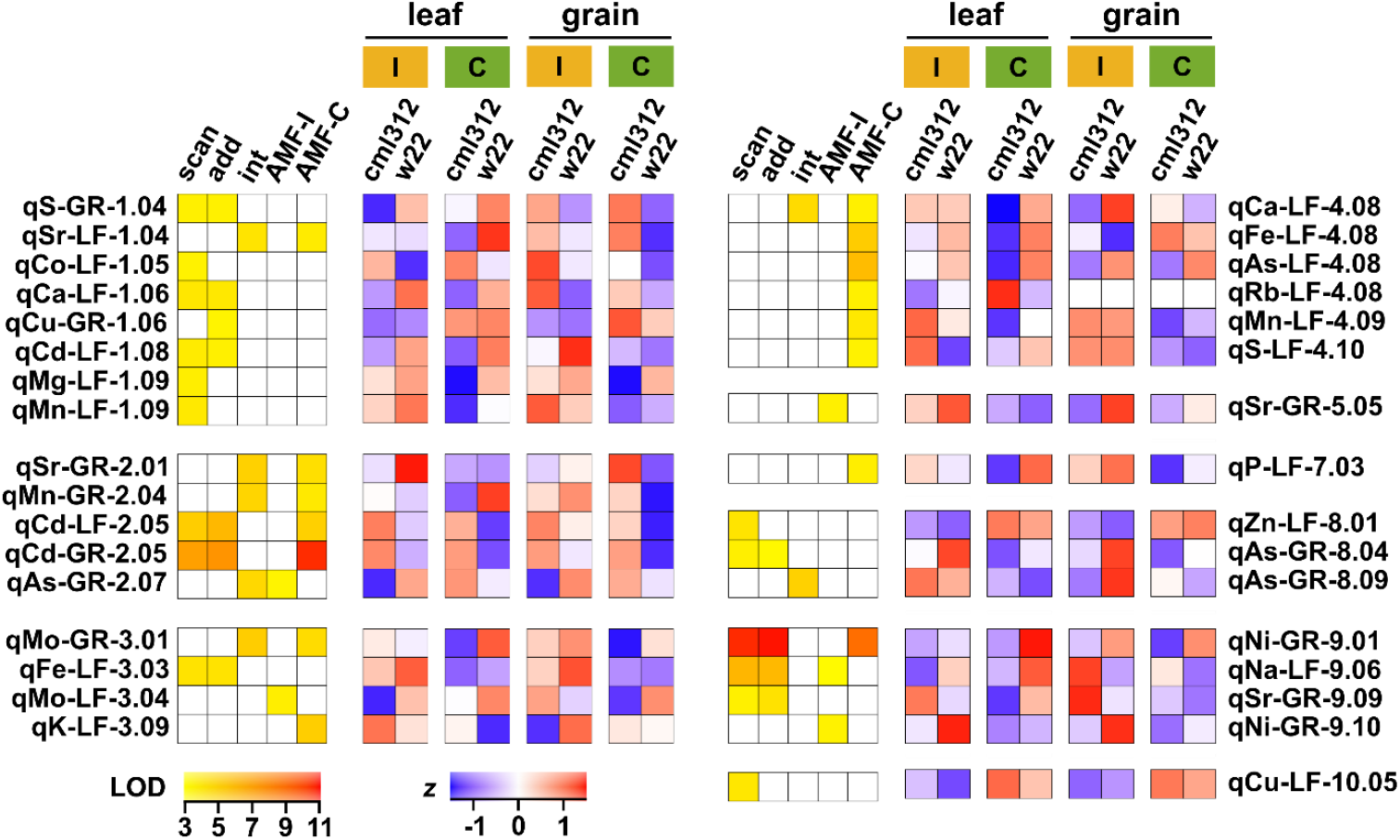
The behaviour of element QTL is contingent on AM status. QTL support (LOD) and allele effects (standardised *z* score for W22 or CML312 allele) for significant element QTL. QTL are named for the associated ion, the tissue type (LF for leaf or GR for grain) and the genetic position (chromosome and bin). LOD support is only shown for the models for which the QTL was significant. Models given as *scan* - all families in a single QTL scan; *add* - AMF status as an additive covariate; *int* - AMF status as an interactive covariate; *AMF-I* - separate analysis of AMF-I families; *AMF-C* - separate analysis of AMF-C families. Effect estimates are shown for both leaf and grain, and both AMF-I (I, yellow) and AMF-C (C, green) families irrespective of whether the QTL was significant in any given tissue/AMF status combination. Effects were standardised separately for leaf and grain.

### Loci with AMF × QTL effects co-localized with QTL showing G×E in a previous multisite ionome experiment

To obtain more rigorous support for our QTL and assess the potential for AMF to impact ionome variation across environments, we compared our findings to a published maize multisite ionome dataset (Asaro *et al*., 2016). This previous work presented grain ionome data for a biparental maize mapping population (hereafter, the IBM data), evaluated during the period of 2005 to 2012, in five different US states, although without specific characterization of the soil microbial community. We re-analysed these data to facilitate comparison with our results and considered each location separately to investigate overlap between AMF × QTL effects in our work and G × E effects in the published study. In addition to QTL identified for individual elements in the previous section, we also included QTL identified for element interaction patterns, including element ratios and the first five latent variables of Factor Analysis, for comparison. We obtained support for 12 of our QTL from the previous data through common signals associated with the same elements in shared genomic bins (Fig. 3A). Interestingly, 10 of the QTL supported by the IBM grain data were identified by us in leaves. The greater power and resolution of the IBM data set increased confidence in our QTL and allowed a more precise estimation of our genomic location (Supplementary Table S1). Nine of the overlapping QTL were detected preferentially in AMF-C plants in our experiments, suggesting AMF communities to be active in the IBM field sites (Fig. 3B). These common QTL also showed evidence of G × E interaction in the previous study (Fig. 3B). Although difficult to substantiate without further data, we speculate that variation in the AMF community between field sites contributed to G × E effects seen in the IBM. For example, strongly supported AMF-C conditional QTL linked to Cd and Ni on chromosomes 2 and 9, respectively, were clearly recovered in IBM data from North Carolina and New York but not detected at all in Missouri (Fig. 3B). Absolute element concentrations indicate that these elements were present in the Missouri environment, although no QTL were detected. Furthermore, the Missouri location lacked signal for an AMF conditional Fe QTL on chromosome 4 that was mildly supported by the other sites, instead showing strong location-specific signals on other chromosomes.

**Figure 3.**
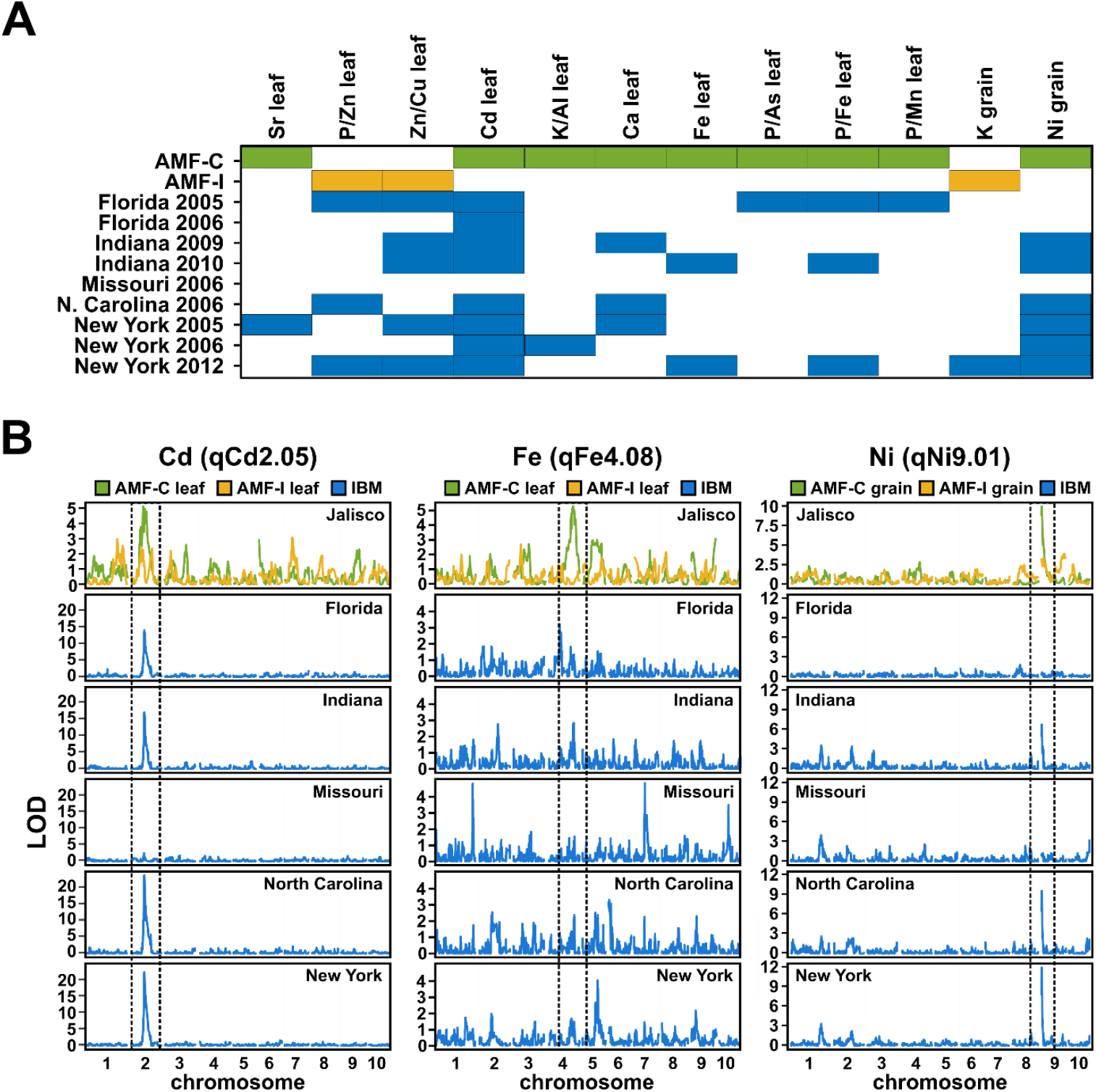
QTL specific to AMF-C families co-localize with element QTL showing G × E in a previously reported multisite evaluation. A) Overlap between element QTL identified in the current AMF experiment and evaluation of the Intermated B73·Mo17 (IBM) mapping population across multiple sites and years (Asaro *et al*., 2016). Coloured boxes indicate a QTL was detected in AMF-C families (green), AMF-I families (yellow) or IBM (blue). B) Genome wide QTL support (LOD) associated with concentrations of the named elements in the AMF and IBM data. IBM plots were averaged over years when multiple data were available. Dashed vertical lines highlight regions containing the named QTL identified in the AMF-C population.

### Patterns in the ionome were correlated with the extent of arbuscular mycorrhizal colonisation

Having observed clear differences between AMF-C and AMF-I families, we proceeded to assess whether ionome variation *within* the AMF-C family was a reflection of differences in the extent of mycorrhizal colonisation. We used microscopy to quantify mycorrhizal hyphae, arbuscules and vesicles from field sampled AMF-C roots in terms of percent root length colonised (as reported previously, AMF-I families were free from root-internal fungal structures (Ramírez-Flores *et al*., 2020)). We observed appreciable colonisation in field sampled roots (hyphae = 23.6% ± 2%, arbuscules = 16% ± 1.4%, and vesicle = 3.4% ± 0.4%), with individual families ranging from 0 to 74.4% in hyphae colonisation (Fig. 4A). To investigate the relationship between the extent of colonisation and the ionome, we used the Boruta feature-reduction method to identify correlations between individual elements and fungal structures. Leaf and grain element concentrations were included simultaneously in a single analysis. The most informative elements were leaf Fe for hyphae, and leaf Fe and Rb for arbuscules (Fig. 4B; Supplementary Fig. S3). Informative elements were used to build Random Forest (RF) models. The trained RF models were used to predict the colonisation phenotypes in the training and whole dataset, and the performance of the models was assessed by estimating the testing (R^2^_T_) and whole dataset (R^2^_WD_) metrics between the observed and predicted data (Fig. 4C; Supplementary Fig. S3, S4). The best performing model was for hyphae (R^2^_T_ = 0.11, p_T_ = 0.17; R^2^_WD_ = 0.29, p_WD_ < 0.001), followed by arbuscules (R^2^_T_ = 0.03, p_T_ = 0.49; R^2^_WD_ = 0.44, p_WD_ < 0.001), and vesicles (R^2^_T_ = 0.041, p_T_ = 0.49; R^2^_WD_ = 0.21; p_WD_ < 0.001).

**Figure 4.**
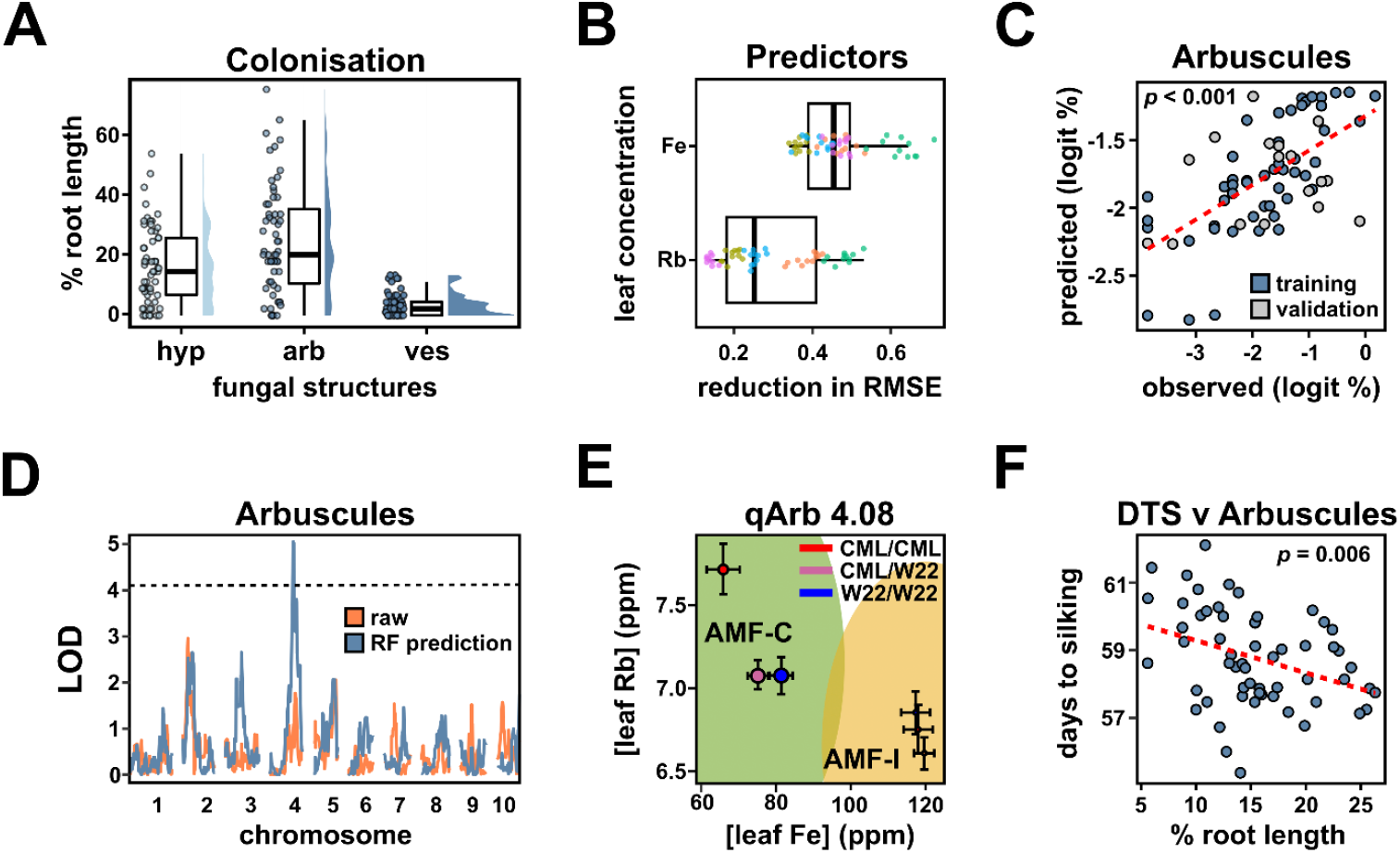
The ionome reflects arbuscular mycorrhizal colonisation. A) Distribution of fungal colonisation across the AMF-C families as percentage root length containing fungal hyphae (hyp), arbuscules (arb) or vesicles (ves). B) Agnostic variable importance for leaf Fe and Rb concentration in the Random Forest (RF) model. The greater the reduction in root mean square error (RMSE) the greater the importance of the variable. C) Correlation between observed and RF model predicted values of logit transformed arbuscule abundance. Observations in the training subset were coloured blue, and data in the validation subset was coloured grey. Validation set r^2^ = 0.041 (p = 0.42); complete set r^2^ = 0.39 (p < 0.001). D) Genome wide QTL support (LOD) support for QTL associated with raw (orange) or predicted (blue) logit transformed arbuscule abundance. The black horizontal line represents the LOD significance threshold (α = 0.1) for identifying qArb_4.08. E) Effect of the genotype at qArb_4.08 on leaf Rb and Fe concentration for AMF-C (green) and AMF-I (yellow) families. Colored ellipses contain 90% of the corresponding families. Points show the average concentration for each genotype, and error bars represent ±1 standard error. The diameter of the point is proportional to the predicted logit transformed arbuscule abundance. F) Scatterplot of Days to Silking (DTS) and predicted logit transformed arbuscule abundance; a mild negative and significant correlation was observed (r = −0.36, p = 0.006).

Our models indicated a relationship between the abundance of fungal structures and the ionome, suggesting that, for a specific environment, a locally trained model could estimate colonisation from element quantification without the need to directly examine roots. We reasoned that ionome predicted values based on a field-wide model might be less biased at the level of the individual than direct observation, given the difficulties of estimation when sampling from a large, heterogeneously colonised root system (Montero *et al*., 2019). To explore this idea, we compared the results of QTL mapping using observed or predicted values of colonisation. Using directly observed values we did not detect any significant QTL (α = 0.1), which was not too surprising given the small size of our mapping population and the difficulty of estimating the trait values. Using predicted values, however, we could detect QTL linked to abundance of arbuscules (Chr 4. Fig 4D) and hyphae (Chr 4. Supplementary dataset). The arbuscule QTL qArb4.08 co-localized with QTL for leaf Rb and Fe concentration - the two ions contributing to the predictive model. The CML312 allele at qArb4.08 was associated with relatively high levels of leaf Rb, low levels of leaf Fe, and fewer arbuscules; conversely, the W22 allele at qArb4.08 was associated with relatively low levels of Rb, high levels of Fe and greater arbuscule abundance (Fig. 4E). The effects of qFe_lf_4.08 and qRb_lf_4.08 QTL were conditional on AM symbiosis and no significant differences were seen between alleles in AMF-I families (Fig. 4E). The conditionality of qFe_lf_4.08 and qRb_lf_4.08 is consistent with their capturing an aspect of mycorrhizal function also reflected in arbuscule abundance. We compared direct observation and predicted values of colonisation in their relationship to plant phenological traits and yield components. Consistent with previous reports (*e*.*g*., (Sawers *et al*., 2017)), we did not find any simple correlation between the abundance of root internal fungal structures and yield components. We did observe greater arbuscule abundance to be correlated with accelerated flowering (Fig. 4F). Similar to the QTL analysis, this correlation was more significant with the predicted arbuscle values than the direct observations.

### Mycorrhizal colonisation promoted alignment of biofortification breeding targets

To understand the potential impact of AMF on host evolvability towards biofortification breeding targets, we compared how well AMF-C and AMF-I families align to a trajectory maximising grain Zn and Fe concentrations, the two mineral elements most commonly lacking in human diets (White & Broadley, 2009). We defined a family to be better aligned if the angle between the major axis of the genetic variance-covariance matrices (G-matrix) and the biofortification target vector was smaller (Fig. 5A; (Noble et al., 2019)). By this criterion, AMF-C families are better aligned with the biofortification target than AMF-I (Fig. 5B). Although AMF-I families showed a higher mean grain Fe concentration (27.5 ppm) than AMF-C families (20.5 ppm), the high Fe concentration was negatively associated with grain Zn concentration (Fig. 5A). Whereas, AMF-C showed a significantly higher grain Zn concentration (19.6 ppm) than AMF-I (13.1 ppm), and the association between Zn and Fe is positive, contributing to its better alignment with biofortification targets. This behaviour was reflected in the AMF-C conditional QTL qGF1 2.10 that was associated with grain factor1 (GF1) that captured variation in both Zn and Fe (Supplementary Fig. S2). In AMF-C families, plants homozygous for the CML312 allele at qGF1 2.10 had greater concentrations of both Zn and Fe in the grain than plants homozygous for the W22 allele, such that selection for the CML312 allele would align with biofortification targets (Fig. 5C). In AMF-I families qGF1 2.10 had no significant effect on Zn or Fe concentrations.

**Figure 5.**
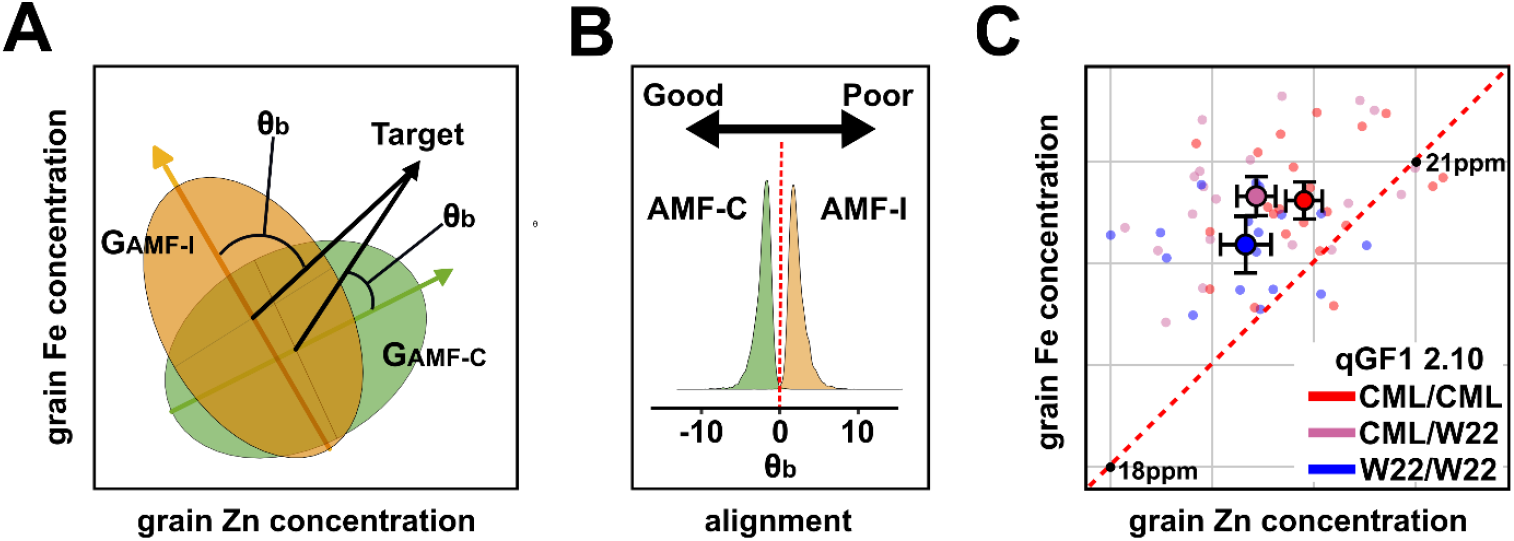
Arbuscular mycorrhizal association may facilitate progress towards biofortification breeding targets. A) The alignment (θb) between the breeding target vector and the major axis of genetic variation in AMF-C (green) and AMF-I families (yellow). B) Distributions showing the effect sizes and corresponding sampling variance of the alignment (θb) of AMF-C and AMF-I families from 5000 Monte Carlo simulations. The negative value indicates that the major axis of genetic variation is well aligned with the breeding target, while the positive value indicates a poor alignment. The x-axis is on a transformed scale: −8 indicates the alignment is approaching 0°, and 8 indicates the alignment is approaching 90°. C) Effect of the genotype at qGF1_2.1 on grain Zn and Fe concentration in AMF-C families. Large points show the mean concentration ±1 standard error. Small points show individual families. The red dashed line indicates equivalence of Zn and Fe concentration.

## DISCUSSION

We have characterised the impact of AMF on mineral nutrition in field grown maize. Using a custom mapping resource comprising both AMF-C and AMF-I maize families (hereafter, mycorrhizal and non-mycorrhizal), we estimated the effect of AMF on the concentrations of 20 elements in the leaves and grain, evaluated the impact of AMF on covariance among elements, and characterised the host genetic architecture of element accumulation with respect to the presence or absence of AMF.

Although the benefit of AMF in cultivated fields has been questioned (Ryan & Graham, 2002, 2018), our data suggest that in our trial AMF improved the uptake of essential nutrients in the field-grown maize. The concentrations of Zn and Cu were significantly higher in mycorrhizal maize families than in non-mycorrhizal families, in both leaf and grain, consistent with previous studies (Lehmann et al., 2014; Lehmann & Rillig, 2015). It has been reported that higher grain yield under P fertilisation reduces Zn concentration through a dilution effect (Zhang *et al*., 2021). Our results showed that mycorrhizal plants maintained greater Zn concentration even though total grain weight per plant was also higher than that of non-mycorrhizal plants (Ramírez-Flores *et al*., 2020). In parallel with an increase in essential micronutrients, mycorrhizal plants showed a significant reduction in the concentration of potentially toxic elements, including Ni and Cd in grain and As and Mn in leaf and grain, again consistent with previous greenhouse studies (Göhre & Paszkowski, 2006; Lehmann & Rillig, 2015; Neidhardt, 2021). As in the pentavalent form of arsenate (AsO_4_^3-^) is chemically similar to inorganic phosphate (PO_4_^3-^) and the two can compete for transport through the P uptake system (Meharg & Hartley-Whitaker, 2002). AMF have been reported to reduce As uptake by down-regulating transporters involved in the direct P uptake pathway (Christophersen *et al*., 2009; Li *et al*., 2018a). We found that a significant correlation between P and As in the leaves of non-colonised plants was no longer significant in mycorrhizal plants, supporting an effect of AMF in changing the As-P relationship. More generally, we found mycorrhizal colonisation altered patterns of covariance among elements, broadly supporting previous greenhouse studies (Gerlach *et al*., 2015; Ramírez-Flores *et al*., 2017), although some specific patterns differed.

Alongside the desirable effects described above, we saw a potentially deleterious reduction in the concentration of the essential micronutrient Fe in both the leaves and grain of mycorrhizal plants. In addition to being essential for plants, Fe deficiency is a widespread problem in human diets making increasing Fe a major goal of biofortification programs (Maqbool & Beshir, 2019). Fe reduction in the shoots of different greenhouse grown mycorrhizal plant species has been reported previously (Tran *et al*., 2019), although AMF have also been reported to specifically increase Fe concentration in the roots (Watts-Williams & Cavagnaro, 2014), suggesting an impact of AMF on root-shoot translocation (Ibiang *et al*., 2017; Xie *et al*., 2019). The interpretation of AMF effects on Fe is further complicated by interactions with Zn status (Ibiang *et al*., 2017). We observed a negative correlation between Zn and Fe in both the leaves and grain of non-mycorrhizal maize families, in line with previously reported antagonism between Zn and Fe nutrition in plant shoots (Ibiang *et al*., 2017). However, this negative relationship between Zn and Fe became positive in the grain of mycorrhizal plants being Zn a major biofortification target (Maqbool & Beshir, 2019). Our analysis of G matrix alignment indicated that although the mean grain Fe concentration was higher in non-mycorrhizal plants, the positive correlation between Zn and Fe in the mycorrhizal plants would promote simultaneous breeding gains towards an increase in the two elements. Although our results were obtained for a biallelic population in a single location, they illustrate a potentially important aspect of the impact of AMF on mineral nutrition beyond increasing or decreasing concentrations in any given genetic background.

We identified 20 element QTLs that showed evidence of interaction with AM status. All AMF × QTL effects were conditional, *i*.*e*., these QTL were specific to either mycorrhizal or non-mycorrhizal plants, although limited statistical power may have prevented identification of more complex genetic architectures. Nutrient acquisition by direct plant uptake or via mycorrhizae follows mechanistically and physiologically distinct pathways (Bucher, 2007; Chiu & Paszkowski, 2019). The contributions of the plant and mycorrhizal uptake pathways are not simply additive but can be better thought of as alternative strategies, one or the other dominating under given conditions (Smith *et al*., 2003). Molecular analyses have supported this view through the identification of diversified families of plant nutrient transporters containing members that play specific roles in direct or mycorrhizal uptake and are regulated accordingly (Casieri *et al*., 2012; Yang *et al*., 2012; Koegel *et al*., 2013; Li *et* al., 2018b; Hui *et al*., 2022). Our identification of conditional QTL is consistent with variation in such pathway-specific genetic components, the resulting impact being expressed preferentially in mycorrhizal or non-mycorrhizal plants. We would hypothesise that typically the balance between direct and mycorrhizal uptake differs spatially across the root system and temporally over the lifetime of the plant, as well as with respect to host genotype. In addition to delivery of nutrients to the arbuscule, AMF secondarily impacts nutrient uptake through enhancement of root growth and modulation of root development (Ramírez-Flores *et al*., 2019). As such, AM symbiosis has the capacity to both mask or exaggerate variation in direct nutrient uptake among host genotypes, a further potential contribution to AMF × QTL effects. For example, our study identified an AMF-specific QTL for Cd in leaf and grain (qCd_lf/gr_2.05). This QTL co-localized with a major locus for maize grain Cd accumulation in a previous study that has been identified as encoding a heavy metal transporter (ZmHMA3) (Tang *et al*., 2021; Chen *et al*., 2022). AMF have been reported to up-regulate the expression of the orthologous transporter gene in rice, and thus promote sequestration of Cd to vacuoles of root cells and reduce the translocation of Cd from roots to shoots (Chen *et al*., 2019; Zhu *et* al., 2022). AMF may also contribute to Cd tolerance in maize through influencing the function of important host Cd transporters, although our results suggest the efficacy of this protection is contingent on the host genetic background.

Root internal colonisation is widely used to characterise the strength of the AMF-host relationship. However, quantifying colonisation typically involves destructive root harvest, staining, and microscopic examination, making it both labour intensive and time consuming. Variation temporally over development and spatially over the root system also make estimation difficult, especially when small, localised samples are collected from the large, and largely inaccessible, root systems of field grown plants. As such, any molecular, metabolic or other indicators that predict colonisation but can be quantified from aerial parts of the plants are of great potential utility (*e*.*g*., the foliar blumenols described by Wang *et al*., 2018). Our analysis suggests that components of the leaf and grain ionome have the capacity to predict AM colonisation. Although we anticipate that any model would have to be trained on a per field/year/population basis, and therefore necessitate some direct root observation, there would still be a great saving of time and effort, especially for large scale field studies. Our models explained, in general, significant variation in the whole dataset (R^2^_WD_), but performed poorly when evaluated in the testing dataset (R^2^_T_), a behaviour consistent with an overfitted model (Ying 2019). This might be the result of a limited dataset consisting of 75 mycorrhizal families, which is then divided into a training and testing set, coupled with the possible presence of noise in the colonisation trait values due to the difficulties in their evaluation discussed above. We hypothesise that a larger dataset might ease the overfitting effect and lead to more conservative results; nevertheless, the modelling approaches we have used identify the most informative predictors from a larger potential set and, given the data, it would be straightforward to combine metabolic, ionomic or other indicators in a single analysis. Mapping the genetic basis of variation in root colonisation is challenging, and where attempted the most informative markers in a given experiment have only described a small proportion of the total variation (0.82-1.14% in winter wheat (Lehnert *et al*., 2017), 6.5% in maize (Kaeppler *et al*., 2000), 8% in sorghum (Leiser *et al*., 2016), 7-16% in durum wheat (De Vita *et al*., 2018) and 4.5-7.1% in soybean (Pawlowski *et al*., 2020)). To date, perhaps the only clear single gene natural variant effect on colonisation to be characterised is that of the Dongxiang wild rice allele of *OsCERK1* in promoting colonisation with respect to cultivated varieties (Huang *et al*., 2020). These results reflect low heritability, in part potentially a result of the aforementioned difficulties in accurately estimating colonisation from small root samples. We had more success in identifying QTL using our ionome-predicted colonisation values than direct observations, which explain ∼ 32% of the variation in predicted arbuscules and hyphae, indicating that for any given individual estimates based on a field-scale trained ionome model may be more accurate than direct observation of a root sample.

The co-localisation of our AMF conditional element QTL and those identified in a far larger study carried out across multiple years and locations (Asaro *et al*., 2016), albeit without consideration of AMF, strongly supports the signals we observed. Furthermore, our observation of AMF conditionality associated with QTL that showed G × E in the previous work is consistent with the hypothesis that variation in AMF communities is contributing to G × E effects. For example, our data indicate that plant genetic variation in the gene *ZmHma3 (Tang et al*., *2021; Chen et al*., *2022)* located at qCd_2.05 will have the greatest impact on plant Cd concentration in the field when the local AMF community is strong. More generally, the AMF contribution to crop G × E is central to any proposition to breed towards superior AMF interactions. In the absence of a significant contribution of AMF to G × E, an improved local AMF community (however defined or promoted) will benefit all plant varieties equally with no need to consider the plants themselves. If, however, the impact of AMF is conditional of the host genotype (*e*.*g*., (Ramírez-Flores *et al*., 2020)) we cannot suppose that crop varieties selected for yield in, for example, an AMF poor environment, will necessarily obtain the greatest benefit for efforts to improve the AMF community in cultivated fields.

## MATERIALS AND METHODS

### Plant material and experimental design

AMF-compatible and AMF-incompatible maize F2:3 families were developed from the cross between a stock homozygous for the *castor*-2 allele in the W22 background and a subtropical CIMMYT inbred line CML312. The experiment was conducted in the summer of 2019 at the UNISEM experimental station in Ameca, Jalisco, Mexico (20.57, −104.04). The field was fertilised at planting with 250 kg/ha of diammonium phosphate (DAP; 18-46-00 NPK) and again at 40 days after planting with 250 kg/ha of urea (46-00-00, NPK). A total of 73 AMF-C and 64 AMF-I families were planted in 3-row plots in 3 complete blocks. Within blocks, AMF-C and AMF-I families were alternated with the order of the families randomised within each block. Further details of the mapping population and experimental design are presented in (Ramírez-Flores *et al*., 2020).

### Determination of elemental concentration by inductively coupled plasma mass spectrometry

Five flag leaves were collected per plot at flowering and oven dried at 70ºC for 48 hours. After drying, 10 cm from the tip were taken from each flag leaf and were pooled to obtain one sample per plot. For the grain samples, 4 kernels were selected randomly from 1-3 ears per plot. Element concentration was determined as described previously (Ramírez-Flores *et* al., 2017). Briefly, flag leaves and grain samples were analysed by ICP-MS to determine the concentration of 20 elements. Samples were digested in 2.5 mL concentrated nitric acid (AR Select Grade, VWR) with an added internal standard (20 ppb In, BDH Aristar Plus).

Concentration of the elements Al, As, B, Ca, Cd, Co, Cu, Fe, K, Mg, Mn, Mo, Na, Ni, P, Rb, S, Se, Sr and Zn was measured using an Elan 6000 DRC-e mass spectrometer (Perkin-Elmer SCIEX) connected to a PFA microflow nebulizer (Elemental Scientific) and Apex HF desolvator (Elemental Scientific). A control solution was run every tenth sample to correct for machine drift both during a single run and between runs.

### Root sampling and quantification of mycorrhizal colonisation

At the flowering time, one plant per row was excavated in a soil monolith (∼15-20 cm from the stalk and 15 cm to 20 cm in depth) using shovels. Gently, the plant was pulled out and shaked to remove as much soil as possible. Twenty root pieces were randomly sampled from a root system and placed in 50mL Falcon Tubes. For the staining, roots were cleared with 10% potassium hydroxide and heated in an autoclave cycle of 15-20 minutes at 121ºC. Cleared roots were stained with 0.05% trypan blue solution in a mixture of 1:1:1 acetic acid, glycerol and water. To quantify AMF colonisation, 15 stained roots were mounted in a slide. Randomly 135 selected microscope fields were observed, and a modified intersections method was used to determine fungal colonisation (McGonigle et al., 1990).

### Data cleaning and processing

All data preparation and analyses were performed in R 4.0.4 (R Core Team 2022). Data was trimmed to remove outliers per element/tissue using R/ graphics::boxplot default criteria, and were adjusted on a per block basis to a spline fitted model using R/stats::smooth.spline against row number to reduce spatial variation at the subblock scale. Mean differences between element concentrations of AMF-C and AMF-I families in leaves and grain were tested by Wilcoxon test with p-values adjusted based on the number of elements using the FDR adjustment method. To further explore the differences of element accumulation between AMF-C and AMF-I families, mycorrhizal responses (MR) of element accumulation were calculated for each element in leaves and grain using the equation: MR = ((AMF-C - AMF-I)/AMF-I)*100% (Watts-Williams *et al*., 2013).

### Mixed effects linear models

The ionomic data collected was collapsed into a single median value per plot for grain and leaf independently. The dataset contained 450 observations for 66 AMF-I and 75 AMF-C families of the F_2:3_ population in 3 different blocks. For each element, a mixed effects linear model was fitted using the restricted maximum-likelihood method with R/lme4::lmer, such that:

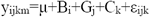

where the response variable y_ijkm_ is a function of the overall mean (μ), fixed effect of the block (B_i_), random effect of genotype (G_j_), and a fixed effect of the AMF status (AMF-I/AMF-C; C_k_), and the residual. Best linear unbiased prediction (BLUP) values for the genotypic effect (G) were extracted using R/lme4::ranef. We calculated fitted values by adding BLUPs to the grand mean for data visualisation and downstream analyses using natural units. Broad-sense heritability for each continuous trait was estimated based on the linear mixed model results using the *bwardr::Cullis_H2* function according to (Cullis *et al*., 2006).

### Multivariate analyses

Fitted values of element B, Na, Al, Se, and Rb showed low variation in either leaf or grain and were removed in the multivariate analyses. To analyse major sources of variation and visualise the element accumulation patterns between AMF-C and AMF-I in leaf and grain, a PCA was conducted using the stats::prcomp function in R. We also calculated the Euclidean distances between centroids of AMF-C and AMF-I families in leaf and grain using the usedist::dist_to_centroids function (Bittinger 2020). Factor analysis was performed to reduce the dimension of element accumulation patterns and extract underlying latent variables that share common trends in differentiating AMF-C and AMF-I. Factor analysis was conducted using the stats::factanal function, with the ‘varimax’ rotation method. The first five factors for grain and leaf were used in the QTL mapping analysis. Detailed loading and scores from the five factors are included in the supplementary dataset.

We calculated pairwise correlations among elements for both AMF-C and AMF-I groups to explore changes in element correlation patterns in response to AMF colonisation. Correlations between elements were performed using stats::cor with spearman correlation method. To test whether correlation matrices are equal between AMF-C and AMF-I, Chi-square test was performed using the decorate::delaneau.test (Hoffman 2021).

Correlations changed in many pairs of elements from AMF-C to AMF-I in our study. We then manually picked pairs of elements that showed large differences in direction or magnitude of correlations between AMF-C and AMF-I families and calculated their ratios and log transformed for further QTL mapping analysis.

### QTL mapping

QTL mapping was performed as described in (Ramírez-Flores *et al*., 2020) using the {qtl} package in R (Broman *et al*., 2003). As phenotypic inputs, the BLUPs for leaf and grain element concentration, ratios, factor scores were used. For mycorrhizal colonisation phenotypes, the median and logit transformed medians were used. Single-QTL standard interval mapping was run using *scanone* function with Haley-Knott regression model and standard parameters. The genotype at *castor* was treated as a covariate (AMF-C = 1; AMF-C = 0; hereafter AMF) for further modelling. To identify any AMF × QTL interaction (Broman & Sen, 2009), four models were considered for mapping: separate analysis on AMF-I (H_0I_) and AMF-C (H_0C_) families, and models considering AMF as an additive (H_a_) or interactive (H_f_) covariate. The LOD significance threshold was established with a permutation test (1000 permutations, α = 0.1). Evidence for AMF × QTL interaction was obtained by comparing H_f_ and H_a_ models, where the difference LOD_i_ = LOD_f_ - LOD_a_ was compared to the threshold difference LOD_thr_i_ = LOD_thr_f_ - LOD_thr_a_ for evidence of possible interaction. Individually detected QTLs were combined into a multiple-QTL model on a per phenotype basis. Where evidence was found of AMF × QTL interaction in the single scan, the interaction was also included in the multiple-QTL model. Multiple-QTL models were evaluated using the *fitqtl* function and non-significant (α = 0.1) terms removed according to the drop-one table.

### Random forest modelling of AMF colonisation levels

A machine learning approach was used to explain and predict the relationship between the ionome and the colonisation level of the AMF-C families using the tidymodels framework (Kuhn & Wickham, 2020). To study the ability of the ionomic data of different tissues to explain the observed colonisation phenotypes, three models with different potential predictors were considered: *ion-leaf*, considering only the grain ionomic variables as predictors; *ion-seed*, considering only the leave ionome variables as predictor; and *ion-everything*, considering both leaf and grain ionome variables and element ratios as predictors. To improve the robustness of the prediction, five different seeds for the random number generator were used (54955149, 100, 22051959, 2500, 161921). The AMF-C subset (n = 75) was split into a training and testing set (75% and 25% of the data, respectively) using the different seed numbers, consisting of five different training sets. The resulting training subsets were used to perform a feature selection process using the {Boruta} package for R (Kursa & Rudnicki 2010) in five bootstrap replicates of the training set. The feature was deemed as relevant if it was selected by the algorithm in at least 20 out of 25 of the seed/bootstrap combinations. The models without features that comply with these characteristics were dropped. Selected features were used as predictors for two different tree ensemble machine learning algorithms: Random Forest modelling was conducted using the {ranger} package (Wright & Ziegler, 2017), and Extreme Gradient Boosting was conducted using the {xgboost} package (Chen & Guestrin, 2016). The number of variables to possibly split at each node, minimal node size and the number of trees for each model/seed/algorithm hyperparameters were optimised independently by cross validation (vfolds = 3, repeats = 10) according to (Kuhn & Silge 2022). The model was finalised with the best combination of hyperparameters and fit with the training set. The trained models were used to predict the values on the training and whole data set, and the average of the prediction of the five different random seeds was considered as the true prediction of the model. The performance of the model was evaluated according to the testing squared r metric.

### Selection trajectory analysis

To understand the impact of AMF colonisation on the genetic architecture of element accumulations and the evolvability to meet future biofortification breeding targets, we applied an evolutionary quantitative genetic approach by quantifying the alignment between genetic variation and the evolutionary trajectory towards maximised biofortification goals. The biofortification goal was defined to obtain maximised grain Zn and Fe as set by HarvestPlus as 38 μg/g dry weight for Zn and 60 μg/g dry weight for Fe (Bouis *et al*., 2011). We extracted genetic variance-covariance matrices (G-matrix) by fitting Generalised Linear Mixed models using Markov Chain Monte Carlo techniques (MCMCglmm) in R. Briefly, the pedigree information was inverted to obtain an inverse sparse matrix, which was then used in the MCMCglmm model with a non-informative prior. The evolvability of crops to meet biofortification targets were determined as the alignment between the main axes of G-matrix and the biofortification target vector (calculated as differences in multivariate trait means from the breeding targets) as described in (Noble *et al*., 2019).

## ACKNOWLEDGEMENT

This study was funded by the Mexican Comision Nacional para el Conocimiento y Uso de la Biodiversidad (CONABIO) project *Impact of native arbuscular mycorrhizal fungi on maize performance* (N ° 62, 2016–2018). RJHS is funded by USDA Hatch Appropriations under Project #PEN04734 and Accession #1021929. GZ and IB were supported by Danforth Center internal funds.

## COMPLETING INTERESTS

The authors declare no completing interests.

## AUTHOR CONTRIBUTIONS

MRRF and RJHS conceived and designed the experiment. MRRF, SPL, BBG, MAMM, GZ, IB contributed to the sample collection and analysis. ML and SPL conducted the data analyses. ML, SPL, MRRF, and RJHS wrote the original draft. All authors contributed to the reviewing and revising of the manuscript.

## DATA AVAILABILITY

Genotype and phenotype data and supplementary dataset are provided on Figshare under the doi’s: https://doi.org/10.6084/m9.figshare.21684947 and https://doi.org/10.6084/m9.figshare.12869867.

## SUPPORTING INFORMATION

**Table S1**. QTLs detected in common with a published multisite ionome analysis.

**Figure S1**. Principal Component Analysis biplots showing element concentration patterns and loadings.

**Figure S2**. Loadings showing the contributions of elements to the first five factors of the Factor Analysis of element contractions in leaf and grain.

**Figure S3**. Random Forest (RF) model results for hyphae abundance.

**Figure S4**. Random Forest (RF) model results for vesicle abundance.

**Dataset S1** Genotype and phenotype data and supplementary dataset are provided on Figshare under the doi’s: https://doi.org/10.6084/m9.figshare.21684947 and https://doi.org/10.6084/m9.figshare.12869867.

## Notes

https://doi.org/10.6084/m9.figshare.12869867

https://doi.org/10.6084/m9.figshare.21684947

### Competing Interest Statement

The authors have declared no competing interest.

