## Supplementary for "Mycorrhizal status impacts the genetic architecture of mineral accumulation in field grown maize (*Zea mays* ssp. *mays* L.)"

**Supplementary Table S1.** QTLs detected in common with a published multisite ionome analysis (Asaro *et al.*, 2016). Significance threshold is alpha = 0.1. QTLs are named by trait and genomic bin, and Chr is the chromosome number. Asaro Trait represents the phenotype in the published multisite ionome analysis (Asaro *et al.*, 2016). Asaro_pos is the genetic position of the QTL for the Asaro Trait in the original analysis in an Intermated B73·Mo17 (IBM) recombinant mapping population (cM). AMF_pos is the position of the AMF QTL in the CML312xW22 genetic map (cM). Est_AMF_pos is the estimated position of the AMF QTL in the IBM genetic map (cM). Asaro_phys _interval represents the 1.5 LOD physical interval of the QTL in the Asaro analysis (MB). For leaf and grain Factor QTLs, the element with an absolute loading value >= 0.5 were considered as causative of the QTL and overlapping with QTLs for Asario traits.

| QTL | Asaro Trait | Chr | AMF_pos | Est_AMF_pos | Asaro_ pos | Asaro_phys_interval |
|---|---|---|---|---|---|---|
| qFe_S_1.04 | Fe | 1 | 60.5 | 59.99 | 82.2 | 256.55 - 265.52 |
| qLF1_1.05 | Mn | 1 | 82.35 | 85.66 | 183.6 | 141.43 - 180.84 |
| qLF1_1.05 | K | 1 | 82.35 | 85.61 | 180.8 | 18.7 - 189.62 |
| qLF1_1.05 | Ni | 1 | 82.35 | 83.31 | 173 | 47.51 - 274.6 |
| qLF1_1.05 | Cu | 1 | 82.35 | 87.12 | 196.1 | 12.98 - 266.74 |
| qCa_1.06 | Ca | 1 | 111.38 | 112.01 | 367.6 | 72.02 - 287.67 |
| qCu_1.06 | Cu | 1 | 112 | 112.01 | 367.8 | 26.83 - 294.16 |
| qLF3_1.07 | S | 1 | 133.5 | 131.33 | 427.7 | 72.71 - 296.97 |
| qZn_Mn_2.04 | Mn | 2 | 46.82 | 45.3 | 46.6 | 14.34 - 20.73 |
| qLF4_2.04 | Cd | 2 | 46.82 | 49.5 | 70.1 | 114.44 - 171.29 |
| qCd_2.05 | Cd | 2 | 58.5 | 58.79 | 214.6 | 155.13 - 164.91 |
| qMo_3.04 | Mo | 3 | 57 | 53.17 | 160.2 | 4.68 - 230.25 |
| qZn_K_4.07 | K | 4 | 96.5 | 98.14 | 308.1 | 56.88 - 227.75 |
| qZn_Mg_4.07 | Zn | 4 | 102.91 | 103.77 | 317.4 | 226.81 - 240.88 |
| qP_Fe_4.08 | Fe | 4 | 112.41 | 87.57 | 287.1 | 185.89 - 233.66 |
| qZn_Mn_5.07 | Zn | 5 | 67.51 | 64.23 | 239.2 | 2.25 - 197.84 |
| qLF3_7.02 | S | 7 | 12.32 | 11.92 | 10.2 | 1.29 - 1.99 |
| qNi_9.01 | Ni | 9 | 1 | 2.05 | 5.4 | 1.33 - 2.26 |
| qSF1_10.04 | Mo | 10 | 40.5 | 43.24 | 88.2 | 142.22 - 146.4 |
| qSF1_10.04 | Fe | 10 | 40.5 | 44.97 | 111.2 | 0.72 - 148.44 |
| qZn_Mg_10.05 | Mg | 10 | 40.76 | 45.11 | 122.4 | 0.72 - 148.44 |

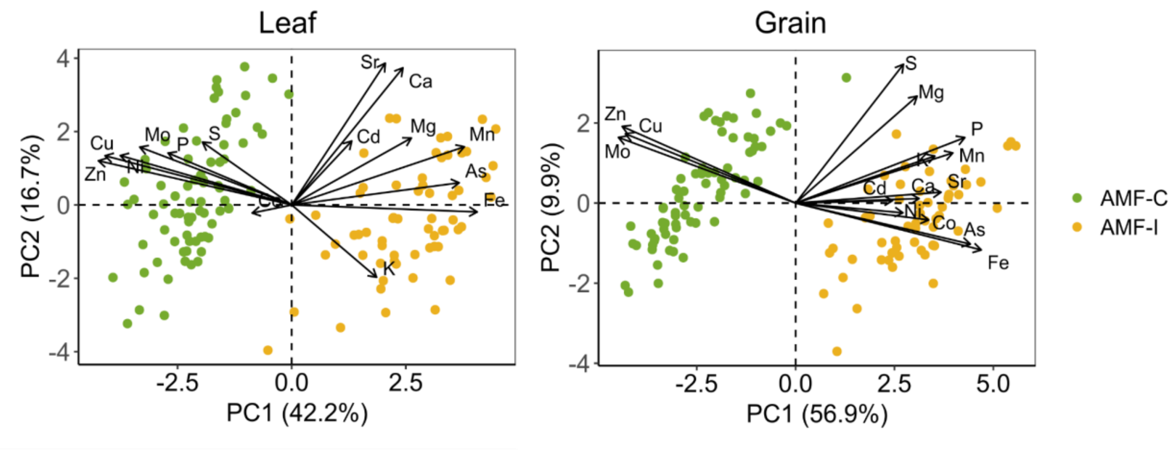


**Supplementary Figure S1:** Principal Component Analysis biplots showing element concentration patterns and loadings of the first two dimensions in leaf and grain of AMF-C (green) and AMF-I (yellow) families.

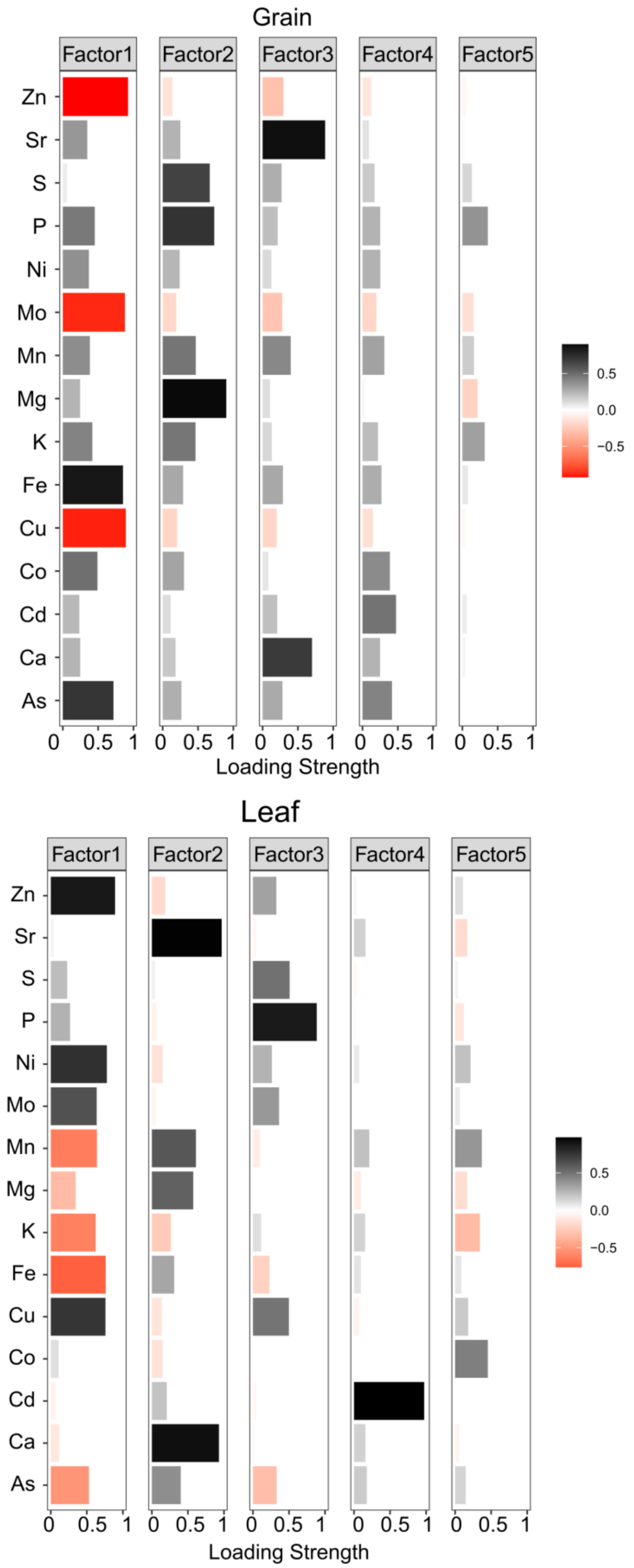


**Supplementary Figure S2**. Loadings showing the contributions of elements to the first five factors of the Factor Analysis of element contractions in leaf and grain.

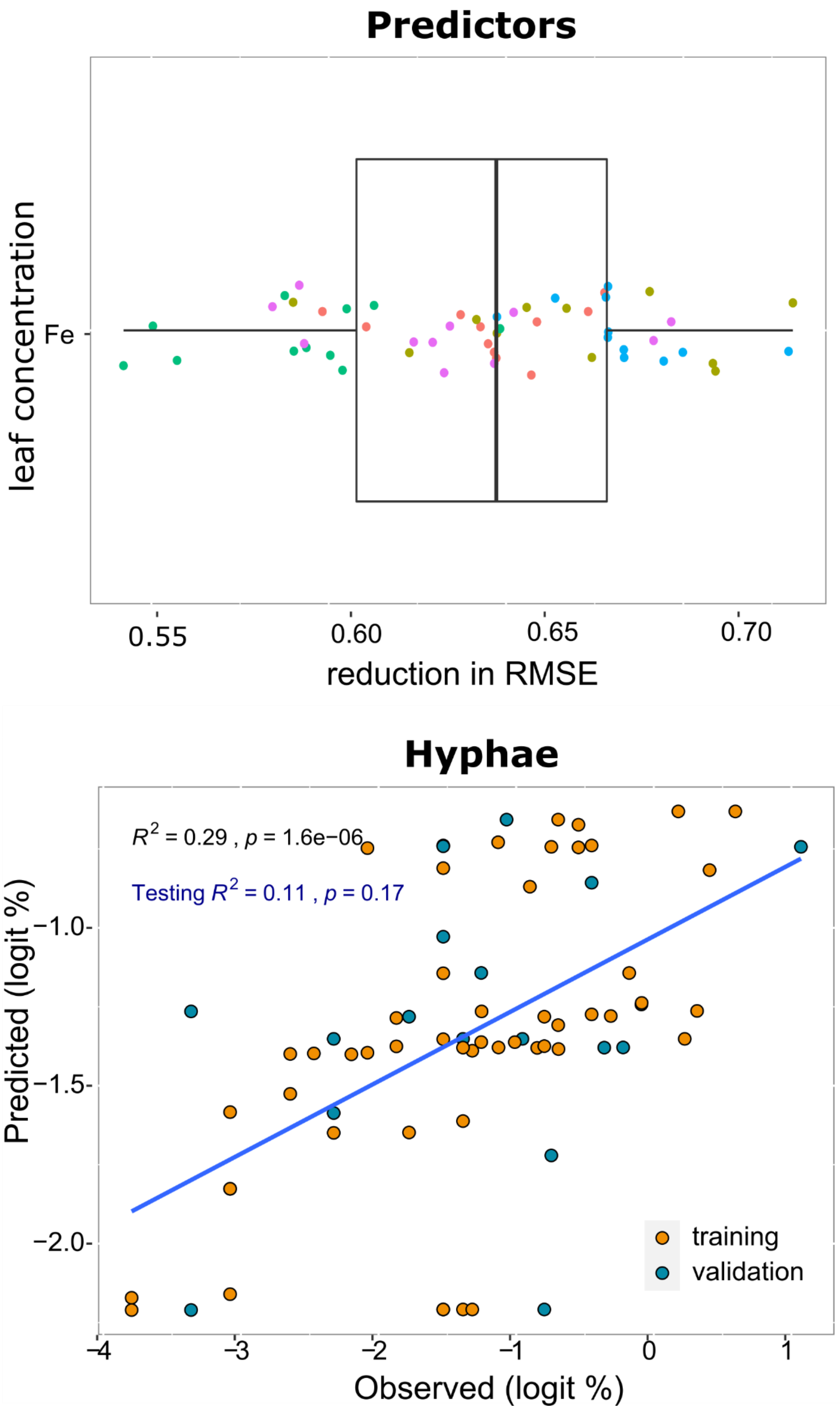


**Supplementary Figure S3**. Random Forest (RF) model results for hyphae abundance. Agnostic variable importance for leaf Fe concentration in the RF model (upper panel). The greater the reduction in root mean square error (RMSE) the greater the importance of the variable. Correlation between observed and RF model predicted values of logit transformed hyphae abundance (bottom panel). Observations in the training subset were coloured yellow, and data in the validation subset was coloured blue.

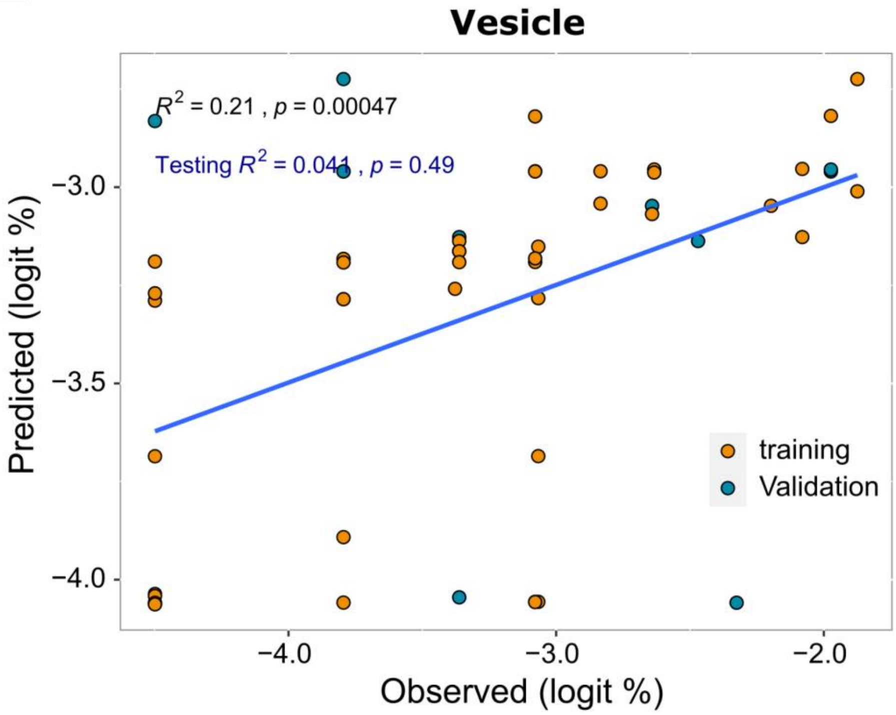


**Supplementary Figure S4.** Random Forest (RF) model results for vesicle abundance. Correlation between observed and RF model predicted values of logit transformed vesicle abundance. Observations in the training subset were coloured yellow, and data in the validation subset was coloured blue.
